# CZ CELL×GENE Discover: A single-cell data platform for scalable exploration, analysis and modeling of aggregated data

**DOI:** 10.1101/2023.10.30.563174

**Authors:** CZI Single-Cell Biology Program, Shibla Abdulla, Brian Aevermann, Pedro Assis, Seve Badajoz, Sidney M. Bell, Emanuele Bezzi, Batuhan Cakir, Jim Chaffer, Signe Chambers, J. Michael Cherry, Tiffany Chi, Jennifer Chien, Leah Dorman, Pablo Garcia-Nieto, Nayib Gloria, Mim Hastie, Daniel Hegeman, Jason Hilton, Timmy Huang, Amanda Infeld, Ana-Maria Istrate, Ivana Jelic, Kuni Katsuya, Yang Joon Kim, Karen Liang, Mike Lin, Maximilian Lombardo, Bailey Marshall, Bruce Martin, Fran McDade, Colin Megill, Nikhil Patel, Alexander Predeus, Brian Raymor, Behnam Robatmili, Dave Rogers, Erica Rutherford, Dana Sadgat, Andrew Shin, Corinn Small, Trent Smith, Prathap Sridharan, Alexander Tarashansky, Norbert Tavares, Harley Thomas, Andrew Tolopko, Meghan Urisko, Joyce Yan, Garabet Yeretssian, Jennifer Zamanian, Arathi Mani, Jonah Cool, Ambrose Carr

## Abstract

Hundreds of millions of single cells have been analyzed to date using high throughput transcriptomic methods, thanks to technological advances driving the increasingly rapid generation of single-cell data. This provides an exciting opportunity for unlocking new insights into health and disease, made possible by meta-analysis that span diverse datasets building on recent advances in large language models and other machine learning approaches. Despite the promise of these and emerging analytical tools for analyzing large amounts of data, a major challenge remains the sheer number of datasets and inconsistent format, data models and accessibility. Many datasets are available via unique portals platforms that often lack interoperability. Here, we present CZ CellxGene Discover (<u>cellxgene.cziscience.com</u>), a data platform that provides curated and interoperable data. This single-cell data resource, available via a free-to-use online data portal, hosts a growing corpus of community contributed data that spans more than 50 million unique cells. Curated, standardized, and associated with consistent cell-level metadata, this collection of interoperable single-cell transcriptomic data is the largest of its kind. A suite of tools and features enables accessibility and reusability of the data via both computational and visual interfaces to allow researchers to rapidly explore individual datasets and perform cross-corpus analysis. This functionality is enabling meta-analyses of tens of millions of cells across studies and tissues and providing global views of human cells at the resolution of single cells.

## Introduction

Cells have been the focus of scientific study for centuries and represent the fundamental unit of life.^1,2,3^ Biology and medicine have long histories of systematically observing, describing, and classifying cells and the anatomical structures that they reside in using assorted methodologies. With each wave of technological innovation comes the discovery of new cell types or states, but also an equally important expansion of knowledge that defines the features of previously described cells in greater detail.

The goal of clarifying the molecular nature of cells has now come within reach due to advances in single-cell measurement technology and concerted community efforts.^4,5^ Over the past 5 to 10 years, research communities have mobilized for projects that span different tissues and organisms, deploying increasingly robust assays and pursuing large-scale characterization of cells that include the Human Cell Atlas (HCA),^4^ Fly Cell Atlas,^6^ Tabula Sapiens,^7^ the Human BioMolecular Atlas Program (HuBMAP),^5^ and many more. These communities have generated a wealth of data that describe how cells vary across organisms, tissues, sex, age, and ancestries.

The incredible progress raises a new challenge, which is that of compiling and distilling the collective value of single-cell data. To unlock the collective value of the work requires addressing pressing barriers to utilizing a large number of massive datasets. Data is often difficult to access and share, with only an estimated 25% of publicly available datasets providing the cell-level metadata needed for reuse.^8^ Even when metadata exists, terms tend to be inconsistent across individual datasets, as do storage formats.

The unique characteristics of single-cell data, including the large number of individual measurements captured compared to other modalities (e.g., microarray, bulk RNA-seq), impose requirements for curation and annotation. These challenges and requirements are not easily met using existing repositories. Several solutions arose due to these unique requirements, offering access to individual datasets and some larger specialized collections, such as Lung Gene Expression Analysis portal,^9^ Allen Brain Map,^10^ and the Single Cell Portal.^11^ Built-for-purpose portals enable rapid publication of studies and dissemination of unique biological features in specific datasets, but lack the scalability and standardization needed for efficient meta-analysis. Even in the presence of such portals, efforts to explore or (re)analyze many datasets must first standardize across individual data portals and resources. Data interoperability is a particularly important challenge for both individual users and for the broader community, to realize the promise of single cell biology both now and in future applications that involve training models or assembling integrated references.

To address the needs for standardized, interoperable, and openly available single-cell matrices, we have developed Chan Zuckerberg CellxGene Discover (pronounced CZ CELL by GENE Discover). This rich platform pairs data and tools that enable scientists to find, download, explore, analyze, and publish standardized single-cell datasets. Open-source and free, it includes contributions from the scientific community and is not confined to a single consortium or funder. It serves as a centralized hub that promotes collaboration across researchers, labs, and consortia. CellxGene is differentiated from other single-cell data portals by its enforcement of a standardized schema for gene, cell, assay, and donor metadata that evolves to address contributor requirements. This standardization provides the foundation for CellxGene’s easy-to-use visual interfaces. In contrast to other repositories that provide only Application Programming Interfaces (APIs) to access datasets individually, CellxGene provides APIs to efficiently query at the level of individual cells across all submitted datasets. CellxGene serves only matrix-formatted data and metadata, which precludes the submission of raw sequence data. This allows for the open sharing of studies that may require controlled access for identifiable sequence data and enables increasingly equitable science for rapid insight into data without the immediate need for data reprocessing. CellxGene Discover builds on the earlier work CellxGene Annotate that enabled exploration, analysis, and annotation of large-scale single-cell datasets, but is optimized for dissemination and reanalysis.^12^

CellxGene is comprised of tools and features designed to address the challenges of compiling and curating data from across a diverse ecosystem by developing scalable infrastructure, visualizations, and interoperability of single-cell transcriptomic data:

1. **Collections & Datasets**: provides the ability to browse or filter individual or collections of datasets and download them with standard metadata annotations in AnnData and Seurat formats.
2. **Explorer**: enables access to single-cell datasets through a no-code user interface so users can execute interactive analyses like differential expression or coloring groups of cells by their metadata.
3. **Gene Expression**: An interactive tool to interrogate gene expression calculated by pooling all datasets for a cell type in any major organ of the human body.
4. **Census**: A client library API in both R and Python, allowing researchers programmatic and efficient access to any custom slice of CellxGene data in a dataset-agnostic fashion.

Together, these features leverage both visual and computational interfaces to allow researchers in the single-cell community and beyond: to explore individual datasets as analyzed and published by the original authors, and to create corpus-wide views, summaries, and meta-analyses of tens of millions of cells across studies. As of October 20th, 2023, CellxGene boasts over 1,102 datasets and 90.2 million cells (56.2 million unique cells), unlocking a new power for any researcher to ask and answer biological questions to clarify off-target effects of drugs across organs, identify of unique marker genes, or interrogate gene expression across cell types for all major human and mouse organs (**Fig. 1**).

**Figure 1.**
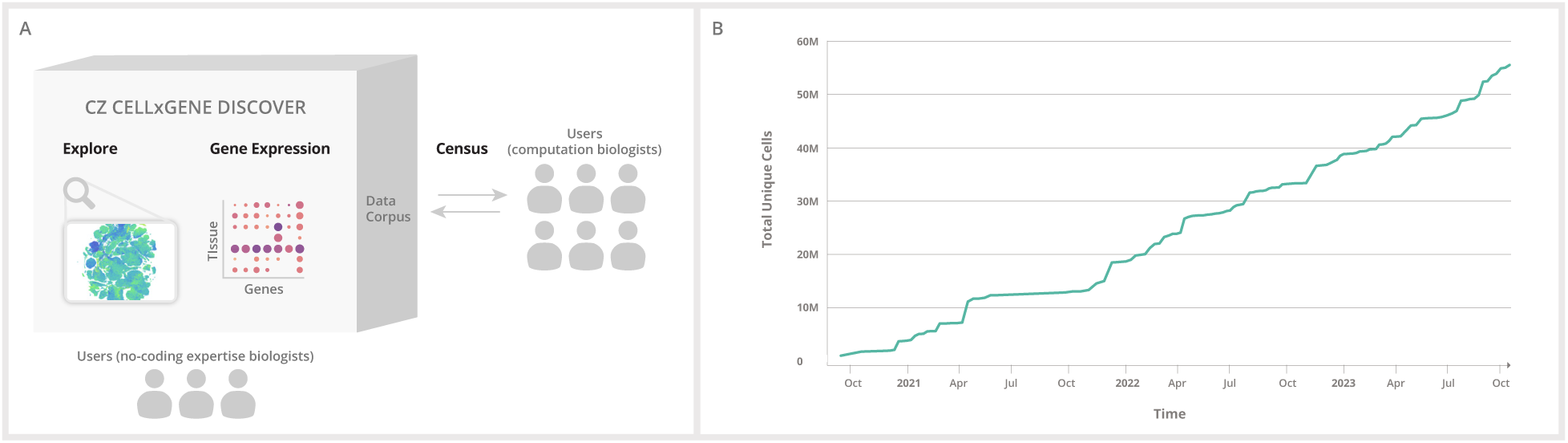
An overview of the CellxGene platform tools and features. (A) The CellxGene data corpus is the central hub, where any dataset can be accessed, downloaded, or explored individually. The data corpus is the foundation for the data visualization and analysis tools, Explorer and Gene Expression, which enable the exploration of individual datasets and cross-corpus views, summaries of gene expression across cells and tissues available in the data portal. A concatenation of all non-spatial transcriptomic data can be accessed via API for downstream analyses through Census. (B) The total number of unique cells available on CellxGene now surpasses 50 million cells.

## Results

### CellxGene data is universally standardized to a minimal data schema

Single-cell transcriptomic datasets from a large number of laboratories, institutions, and consortia are available via websites and portals.^13^ Each portal serves the needs of its dataset, but typically fails to enable collective use. This presents a barrier to downstream use of the data and queries that run across many datasets. To address this challenge, we defined requirements and standards for a minimal schema that reflects widely used terms aimed at enabling corpus-wide searches, filtering of data, and data analyses. Our driving motivation was to enable dynamic integration and queries of all the data or a subset of cells of interest to a biologist.

### A minimal cell-level schema

Metadata is critical for the reuse of data but often inconsistently captured or represented in single-cell datasets. A first key step toward ensuring greater reusability of data was defining a core set of metadata fields and ontologies. We sought to define a schema that targeted data integration. Central to single-cell biology, integration enables the compilation of datasets into atlases.^14,15^ It allows for the inference of multi-modal measurements, cell type prediction, and cross-species analysis.^16^ We collaborated with the single-cell community — including members of Human Tumor Atlas Network (HTAN),^17^ HuBMAP,^5^ and HCA^4^ — to develop standards in line with the key information frequently utilized by meta-analysis studies to properly integrate data or to identify biological variables correlated with gene expression.^18,19,20^ To avoid deterring or inhibiting data submission and adoption, we limit the schema to 11 required fields considered most valuable for data integration and reuse (with additional information permitted at the user’s discretion). The resulting requirements were encoded into a minimal, versioned schema that all submitted data must adhere to and be validated against during the submission process (**Fig. 1, table S1**).

The required fields represent attributes that are often variable within or across studies and are often identified as strong covariates correlated with gene expression variation within cells.^21,22^ Similar to previous experimental data coordination efforts, established ontologies and other community resources are used for standardization wherever possible for consistency and to improve the filtering capabilities of datasets (**table S2**).^23,24,25^

To fully capture gene count information for each dataset, a layer of raw data, meaning non-normalized, is required for submission of data from all transcriptomic assays to fulfill common computational reuse cases.^19,26,27,28^ The corpus-wide availability of raw counts greatly enhances data accessibility by openly serving reusable data products for studies with controlled access sequence data and prevents computational resources from being used towards re-alignment for cases where the original alignment will suffice.

Additionally, CellxGene does not recluster or perform analyses on individual datasets. As a result, a minimum of one two-dimensional embedding (such as UMAP, tSNE, PCA, etc.) is required to facilitate dataset visualization within the Explorer interface (see Section 3: Scalable tools allow biologists to explore, query, and analyze CellxGene data). Importantly, CellxGene allows for multiple embeddings, enabling the sharing and exploration of diverse data representations.

Metadata for a given dataset is not constrained to CellxGene’s minimal schema; it is extensible and allows for the submission of additional metadata that contributors consider valuable. In the case of coordinated efforts or consortia, the system fully supports additional fields that have been standardized across studies to encourage and enable meta-analysis. The full schema specifications have been adopted by multiple consortia, including the HCA,^4^ BRAIN Initiative,^29^ and Kidney Precision Medicine Project (KPMP),^30^ and are available to the research community to support interoperability with current and future efforts (**table S1**).

### Schema evolution

Schema evolution is anticipated given the advancement in data generation techniques and data analysis technology, especially in a rapidly advancing scientific field like single-cell biology. We designed the minimal schema to be supportive of changes and review the data corpus every 6 months for opportunities to make it increasingly comprehensive and standardized. With each schema update, previously submitted data are migrated to meet the new schema, even if that requires re-curation (see Methods). These data migrations ensure that all datasets are consistently described by the most current standards at all times, and users will have consistent filtering and integration experiences independent of when a given dataset was initially submitted. The full CellxGene schema changelog is available (**table S1**).

### Submission workflow

CellxGene welcomes data contributions meeting current submission criteria (**table S1**) from any individual contributor, laboratory, or institution. Upon request from data contributors via email to, a dedicated curation team creates a new private Collection, defined as a group of datasets that are part of a study or publication, with a contributor-provided title, description, contact, associated consortia or projects, as well as any external URLs (**table S2**). To enable collaboration and journal reviewer access without registration, the curation team provides the contributor with a URL to their new Collection. This URL is permanent and will not change when the Collection is made public. The benefit is that contributors do not need to update their manuscript or other text for publication where the URL is referenced. This URL is also obscure such that until the Collection is made public, the URL is only viewable by the contributor and anyone they share the URL with. Once a dataset fulfills the schema requirements, it is uploaded to the Collection as an H5AD file in AnnData format.^31^ Upon upload, the AnnData object is updated with human-readable gene symbols and ontology labels based on the submitted identifiers. A Seurat object is then created from the updated AnnData.^32^ Both formats are made available to consumers for download to enable reuse in a variety of downstream single-cell analysis toolchains. In addition, an internal format for visualization in Explorer is produced, as described in Methods. Gene symbols and ontology labels are mapped consistently throughout the data corpus via a specific ontology release version defined in the schema for each community resource.

Additional details on the data submission process are available on the CellxGene documentation pages (**table S1**).

### Community Resource for Standardized Cell Resolved Measurements

CellxGene hosts the largest curated collection of publicly available single-cell data, with data from 252 tissues and 34 unique assay types. Although the corpus does contain data from samples across 90 diseases and various organisms (e.g. *Mus musculus*, *Macaca mulatta*, *Callithrix jacchus*, and *Sus scrofa domesticus*), the resource has focused on healthy human (**Fig. 3A**). The data schema (**Fig. 2**) provides the ability to assess the comprehensiveness and deficits of CellxGene data. Current data accumulated is insightful for identifying areas of data generation, at both the assay and sample level, that are poised for meta-analysis as well as deficits that need to be addressed in the future.

**Figure 2.**
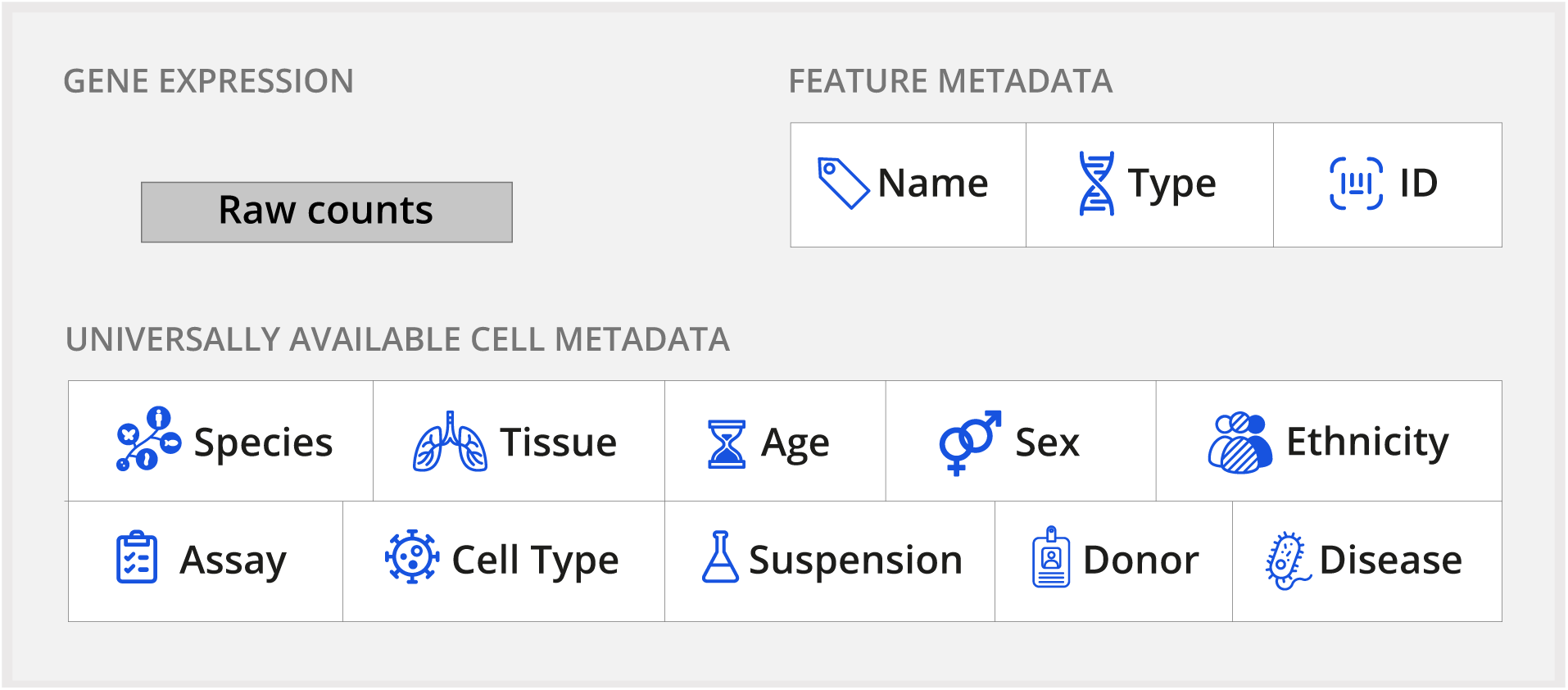
Description of the CellxGene schema. CellxGene requires all data to conform to a standard metadata schema. The schema requires raw counts (e.g. mapped but unnormalized) as part of data submission. Required metadata covers 10 generally available categories that are completed for each sample and cell to increase reusability for downstream analyses. An additional metadata category, not shown in the figure above, is the is_primary_data field. This field is used to mark each observation as “primary” exactly one time throughout the corpus so that cross-corpus aggregations can avoid redundant observations in their analysis.

**Figure 3:**
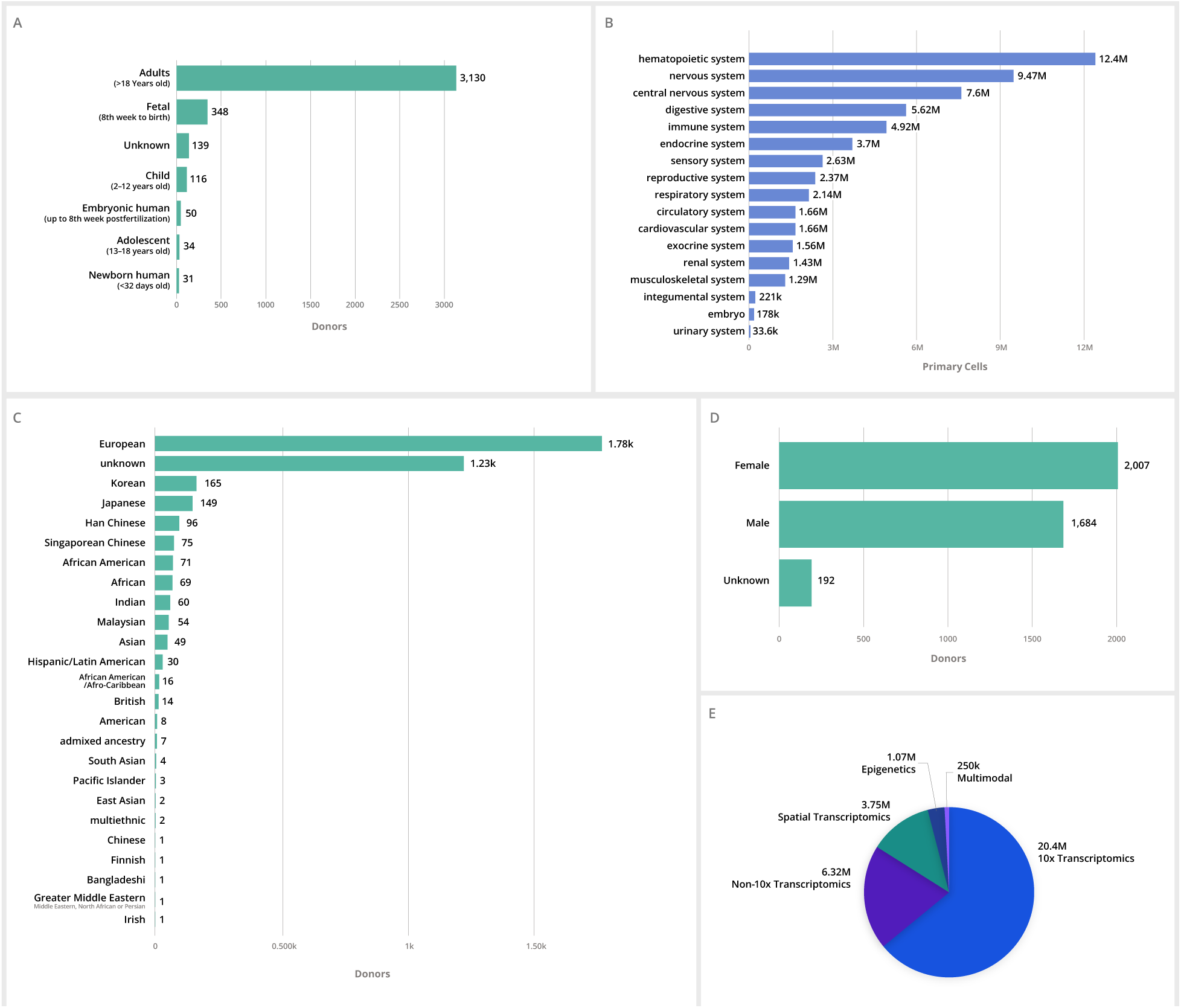
CellxGene data corpus across various metadata categories. (A-D) A breakdown of total unique cells across all major organ systems in CellxGene and total healthy human donors across self-reported ethnicity, developmental stage, and sex available in the CellxGene data portal. (E) The majority of data available in the CellxGene data portal is from 10X Genomics transcriptomics assays, with over 20 million cells available for download and exploration. Additional modalities, including non-10X transcriptomic assays, spatial transcriptomics, epigenomic, and multimodal data types, are supported if they meet the minimal schema requirements.

Of the 50+ million cells available in the data corpus, 84% of the total cells in the corpus are human, 15% are mouse, and less than 0.5% are from other species. Of the human data, approximately 66% is defined as coming from healthy human donors, while the remaining 34% span 90 unique diseases. The number of donors broken down by metadata category, with an appreciable coverage of cells from donors for the hematopoietic (1,890 donors) and respiratory systems (688 donors) (**Fig. 3B**). This accumulation reflects the significant data generated and contributions of single cell methods to the COVID-19 pandemic. However, within these systems, and other systems present in the corpus, diverse ethnicities and age groups are grossly underrepresented (**Fig. 3C**). Samples from female tissue donors are slightly more numerous across the data corpus (**Fig. 3D**).

The CellxGene schema and underlying architecture have been designed for extensibility to new modalities and growth of the data corpus. Current data is primarily single-cell transcriptomic, although CellxGene accepts additional modalities to support the growing number of studies that incorporate a multitude of assays and co-assays (**Fig. 3E**). New data will require ongoing evolution of refined human cell types and tissues in a standardized, structured way by leveraging and contributing to the Cell Ontology (CL),^33^ Human Ancestry Ontology (HANCESTRO),^34^ and UBERON.^35^

CellxGene provides two main ways to interactively browse data: the Collections and Datasets pages. The Collections page (https://cellxgene.cziscience.com/collections) allows users to view all Collections, defined as thematic groups of data, typically grouped by a publication, in the data corpus and filter by metadata categories. If a Collection has a publication DOI, then its author information, journal, and publication date are retrieved from the Crossref service to filter by Publication Date or Publication. Each collection also provides contributor and provenance information including contact, narrative description, and an association with a consortium if appropriate. From the Collections page, one can view high-level attributes of the data contained within each Collection (e.g. organism(s), tissue(s), and disease(s)), and a short citation of the associated publication, if one is present.

The Datasets page (https://CellxGene.cziscience.com/datasets) allows for viewing and filtering all datasets in the data corpus by the same metadata as the Collections page, plus cell and gene counts, and provides access to the downloadable files and Explorer visualization for each dataset. Clicking on a Collection title from either search page will direct users to the corresponding Collection page where one can view the Collection information and a Dataset table that provides summary metadata for each Dataset in the Collection as well as file and visualization access (**fig. S1**).

### Scalable tools allow biologists to explore, query, and analyze CellxGene data

A universal view of atlas data represents an important next step for atlas data to encourage greater insight from the immediate users of the data and wider sections of the scientific community. The preponderance of tissue and multi-organ atlases provide ample data but raise a myriad of challenges related to interoperability, data format, and the ability of computational tools to quickly query 10s or 100s of millions of cells. CellxGene addresses several challenges associated with data aggregation and standardization to enable researchers with wide ranges of expertise to visualize, explore, access, and reuse single-cell data. Currently, we provide three main tools: Explorer, Gene Expression, and Census, all of which utilize standardized data to lower the bar for integration as well as allow immediate exploration of biological insights.

### Explorer allows interactive exploration and analysis of individual single-cell datasets up to 4M cells

The majority of single-cell visualization portals rely on precomputed values that limit interactive features on datasets and generally are tuned to host datasets of 250-500k cells.^36^ Portals have struggled to keep up with the growth of dataset size, now reaching the order of millions of cells per study.^37,38^ As the field scales, and perhaps importantly as more integrated datasets become available, it is critical to have the ability to dynamically visualize and perform basic analysis functions on millions of cells. Explorer, a feature of CellxGene, is a visualization platform that allows researchers to dynamically explore, compute, and query individual datasets for up to 4.3 million cells with minimal latency (see Methods).

With Explorer, researchers can explore gene expression by visualizing individual genes or groups of genes (**Fig 4, inset**); performing differential gene expression; identifying marker genes; and visualizing continuous (e.g., UMI count) and categorical (e.g., cell type) cell metadata.

**Figure 4.**
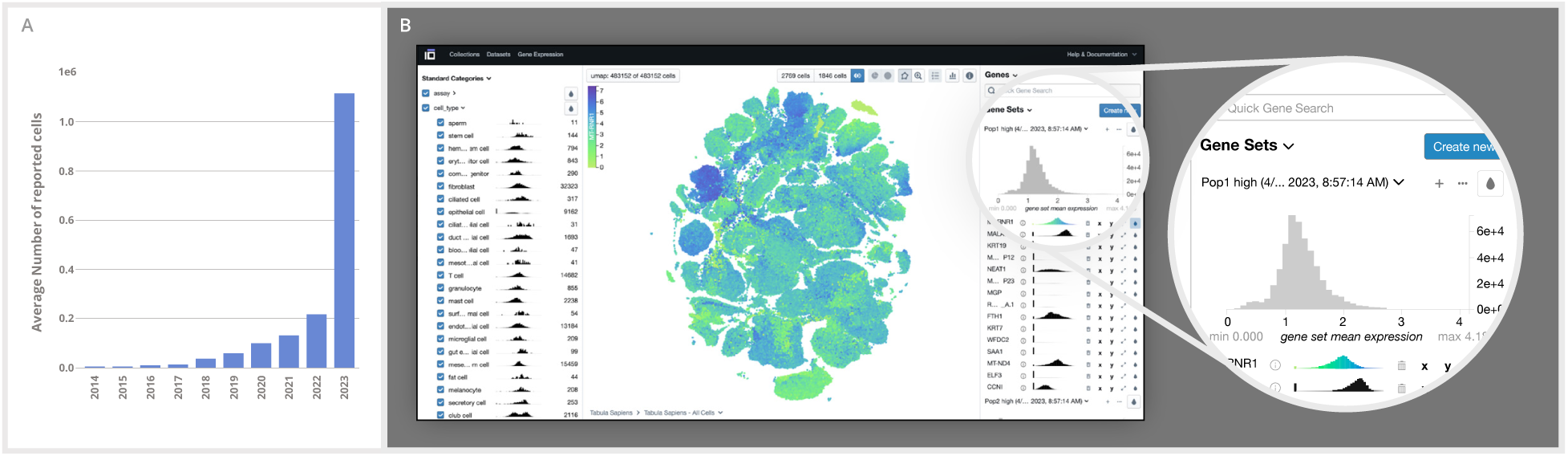
Explorer enables interactive analysis of single-cell datasets. A UMAP of all 483,152 cells in the Tabula Sapiens dataset available on Explorer visualized by the expression of MT-RNR1, a gene that encodes for an rRNA responsible for regulating insulin sensitivity and metabolic homeostasis.^39^ UMAPs can be visualized based on metadata categories, including cell type and other metadata categories, or by the expression of one or multiple genes, as shown above.

In addition, Explorer offers a dynamic interface that allows users to rapidly explore co-variation and trends that are not typically captured or presented in pre-computed interfaces. Users can facet these visualizations based on gene expression, metadata, and the embedding itself. The combinatorial nature of these affordances facilitates arbitrary comparisons through cross-filtering, subsetting, and coloring subpopulations of the data (**Fig. 4**).

### Gene Expression allows gene expression queries across the corpus of data

An immediate opportunity of single-cell atlases is to offer researchers the ability to interrogate the expression of genes across cell types in a specific biological context, such as tissue, disease, ancestry, sex, and developmental stage. To meet this need, we developed Gene Expression, an interactive tool with an intuitive visual interface to explore gene expression across all RNA datasets hosted on CellxGene that meet assay and quality criteria (see Methods).

Gene Expression displays heat maps for genes across cell types and tissues to visualize gene expression. For any given cell type and gene, the dot color represents the average gene expression of all cells annotated accordingly. The size of each dot represents the percentage of cells within the cell type that express that gene (**Fig. 5A**). This visualization reveals high-level differences in expression patterns across cell types. The combination of these metrics in a grid of [*cell types* by *genes*] allows researchers to make qualitative assessments of gene expression between user-defined subsets of cell types and tissues. Researchers can filter by tissue, publication, sex, self-reported ethnicity, and disease to tailor their results to a subset of the data they are interested in or deem most relevant (**fig. S1**). The Group By functionality reports gene expression for cell types stratified by the selected category (i.e., sex, self-reported ethnicity, and disease), further enabling researchers to identify the genes and mechanisms that define and differentiate cell types in different biological contexts (**Fig. 5B**).

**Figure 5.**
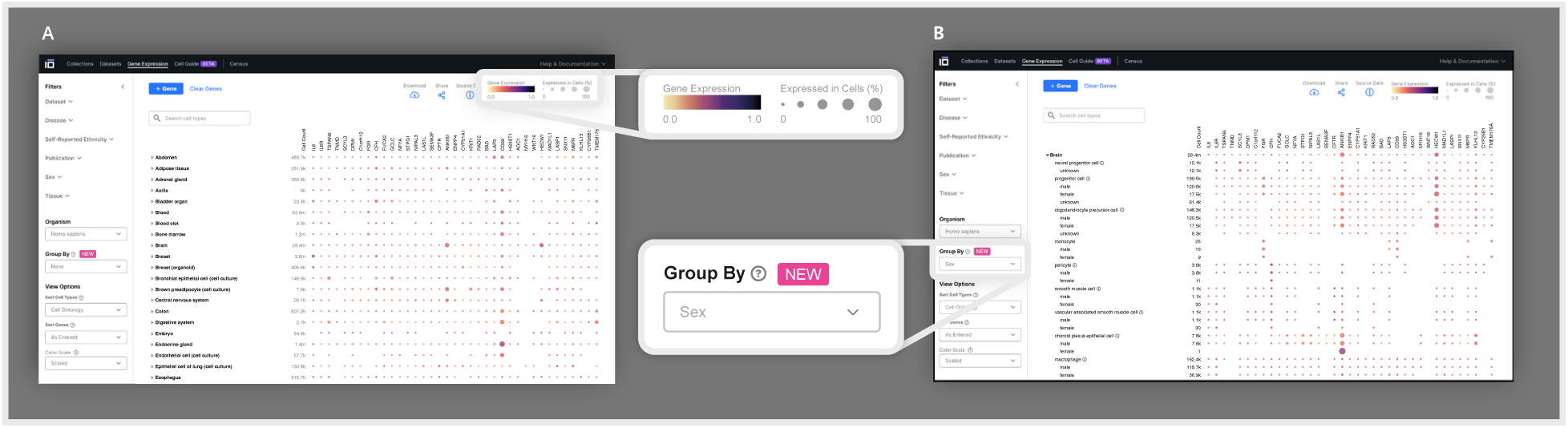
Gene Expression enables the cross-dataset visualization across tissue. (A) A heatmap generated using Gene Expression visualizing the mean gene expression of specific genes across and within all tissues and cell types present in the data corpus. The quantity of cells used for calculating the mean gene expression within each tissue and cell type is indicated in the leftmost column of the heatmap under “cell count.” The gene expression is displayed using two visual elements: color, representing the mean gene expression, and size, signifying the proportion of cells in each cell type or tissue expressing the respective gene. (B) A heatmap generated through the utilization of the Group By feature, demonstrating the variation in gene expression among different cell types according to sex. Group By allows researchers to group mean gene expression values by specific metadata values, including, sex, disease, and ethnicity.

### Normalization of the Gene Expression data enables the core functionality of Gene Expression

Gene Expression’s underlying data is a concatenation of data of cells from transcriptomic sequencing assays that measure gene expression and do not require gene-length normalization (see Methods). It is built off the raw counts (i.e., mapped but unnormalized reads) from each individual dataset; these raw counts undergo a standard pipeline of data processing and normalization (see Methods). Gene Expression data can be downloaded from the CellxGene documentation site (**table S1**).

We chose to normalize raw counts from this dataset to log-transformed scaled pseudocounts (“log transformation”) based on requirements that we established to meet a general use case of enabling scientists to explore gene expression across all eligible datasets in the corpus: it is a minimal, interpretable manipulation of the data; it is scalable to millions of cells; and it stabilizes the variance to a smaller range, suitable for visualization and rapid exploration.

We evaluated the efficacy of log transform normalization in preparing this data for Gene Expression’s primary use cases: (1) displaying the mean gene expression values for each cell type on a comparable numerical scale, and (2) identifying marker genes for each cell type population. Given these focused use cases and technical requirements, the scope of our evaluation was to determine whether log transformation of scales pseudocounts was a reasonable selection, comparable to other common normalization approaches which met our requirements such as quantile normalization, rather than a full investigation of all available normalization methods.

It is important to note that log transformation does not aim to eliminate batch effects, as it merely normalizes the data without full integration. However, we wanted to understand how strongly batch effects might influence *aggregated* (averaged) normalized gene expression values. If log transform normalization effectively mitigates batch effects in the averaged values, then we would expect to see similar average normalized values of gene expression between batches. To test this, we normalized the data, stratified by covariate values, and then computed the mean gene expression vectors for each covariate value (e.g., each dataset, each assay, etc.). To assess the differences between these vectors, we calculated the log(fold change) between each pair (see Methods). We looked at Dataset ID and Sequencing Assay as potential covariates. Differences in mean gene expression between batches are greatest and vary the most in raw counts (*σ_raw_ dataset* = 0.332, *σ_qn_ assay* =0.336), and lower after quantile normalization (*σ_raw_ dataset* = 0.170, *σ_qn_ assay* = 0.168). We can observe a further decrease for values normalized with log transformation (*σ_ln(CPM+1)_ dataset*=0.108, *σ_ln(CPM+1)_ assay*=0.103)(**Fig. 6**). We have also conducted a repeated measures ANOVA analysis to assess the impact of covariate values on the average gene expression values on a number of cell types. Across a set of marker genes and housekeeping genes, our results show that for most cell types, we do not have sufficient evidence to say that there are significant differences between the average normalized gene expression values across covariates. This suggests that batch effects are mitigated compared to raw counts; however, users should be advised that the displayed values may still be affected by batch effects, although this is partially mitigated by normalization and aggregation.

**Figure 6.**
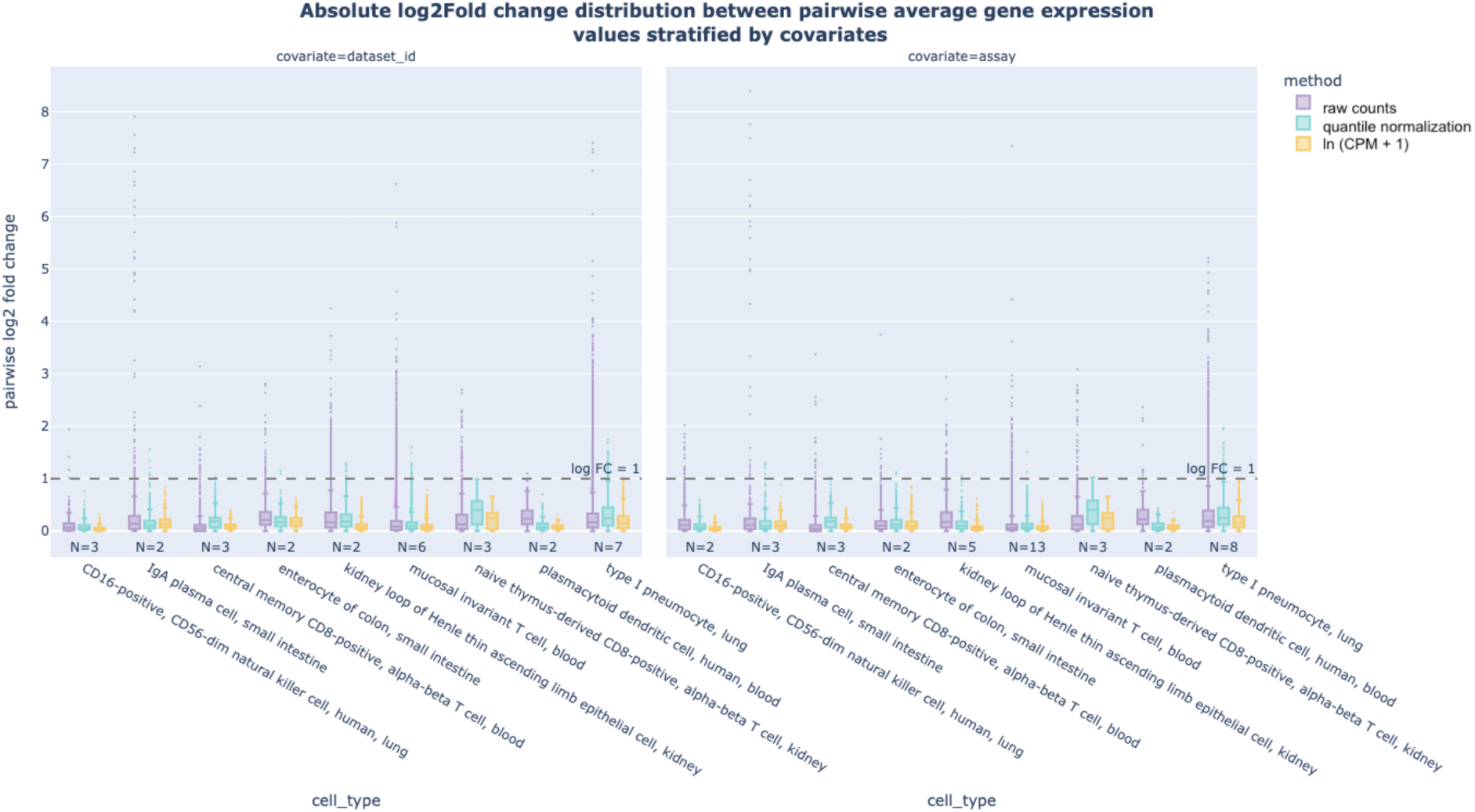
Batch Effects Present in Average Gene Expression Values. Distribution of absolute log fold change values between pairwise average gene expression vectors stratified by covariates. Each point in the distribution represents the log fold change between the average gene expression values of a particular gene between two covariate values (e.g. one point in the Dataset ID covariate plot represents the log fold change between the average gene expression values of a particular gene in two different datasets). N is the number of covariate values (e.g. number of assays or datasets) per cell type. We consider all genes expressed in a particular cell type. Spread of log fold change between batches is highest for raw counts (*σraw dataset = 0.332*, *σraw assay* = 0.336), followed by quantile normalized values (*σqn dataset* = 0.170, *σqn assay* = 0.168), and log transform normalized values (*σln(CPM+1)*dataset = 0.108, *σln (CPM+1)assay* = 0.103).. Visualization is based on a random subsample of the data.

**Figure 7.**
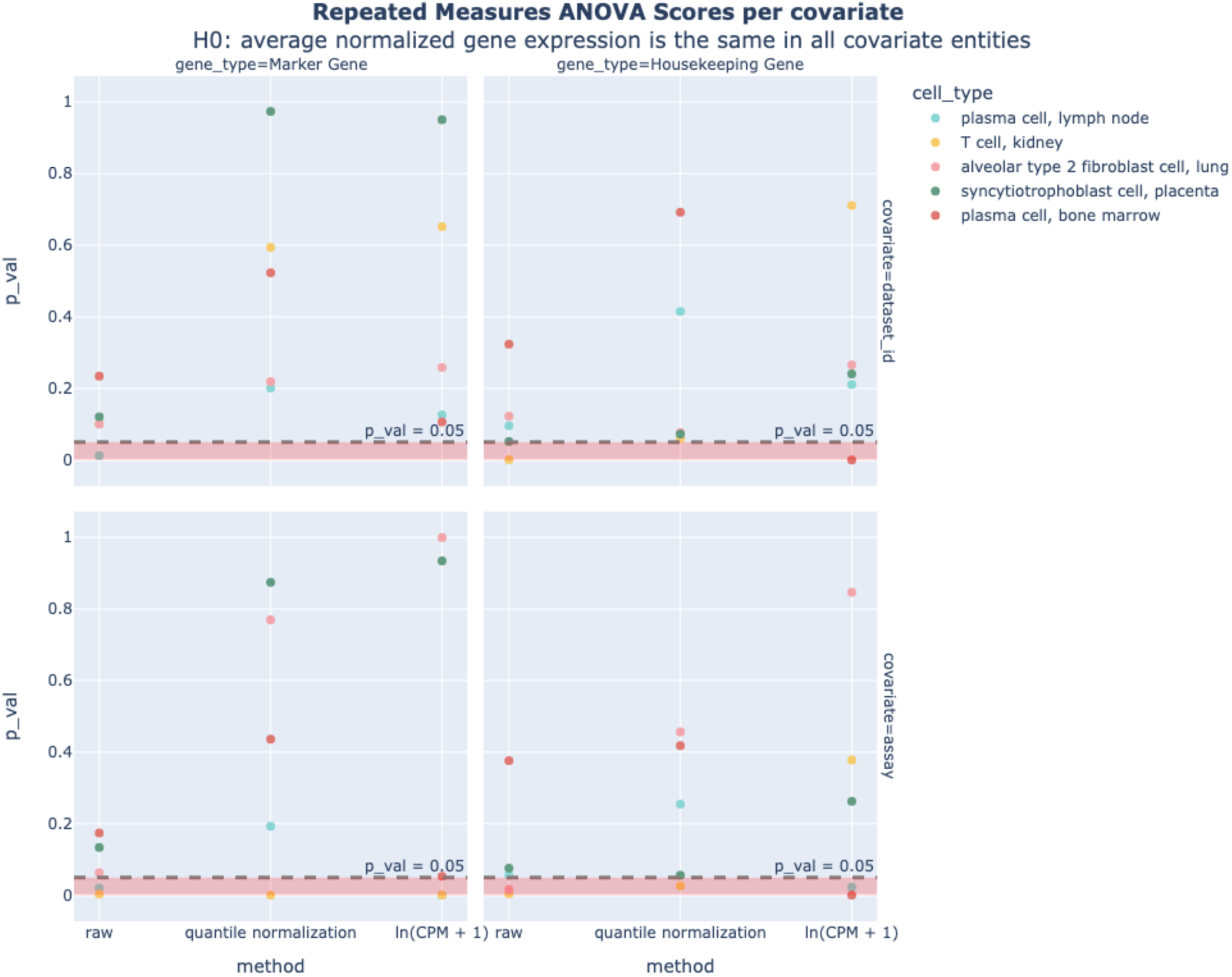
ANOVA on average normalized gene expression values. Results for one-way repeated measures ANOVA scores conducted on marker and housekeeping genes in five different cell types. Results show that for most cell types, we don’t have sufficient evidence to say that there is a statistically significant difference between the average normalized gene expression values among covariate values (9/10 and 6/10 *p* values for ln(CPM+1) for dataset_id and assay > pval = 0.05, aggregated over marker and housekeeping genes)

Next, we analyzed how log transform normalization might influence our ability to recall canonical marker genes. We compared the sensitivity of a *t*-test in recalling marker genes from HuBMAP^5^ for 127 cell types from 14 tissues on quantile-normalized data and log transform normalized data (details in Methods). Cell type selection was based on data availability; where two cell types were collinear in the Cell Ontology, we removed the more basal type. We found comparable performance across normalization methods, with an average sensitivity of 33% for both quantile normalized values and log transform normalized values (**Fig. 8**). The probability of observing this level of sensitivity from the log transform normalized values by chance for each cell type ranges from p=0.00 – 0.04; the moderately low sensitivity values are likely influenced by the fact that the HuBMAP dataset we use as our “ground truth” includes annotations from a variety of different assay types and are quite noisy. Overall, these results indicate that log transform normalization does not detrimentally affect our ability to recall marker genes.

**Figure 8:**
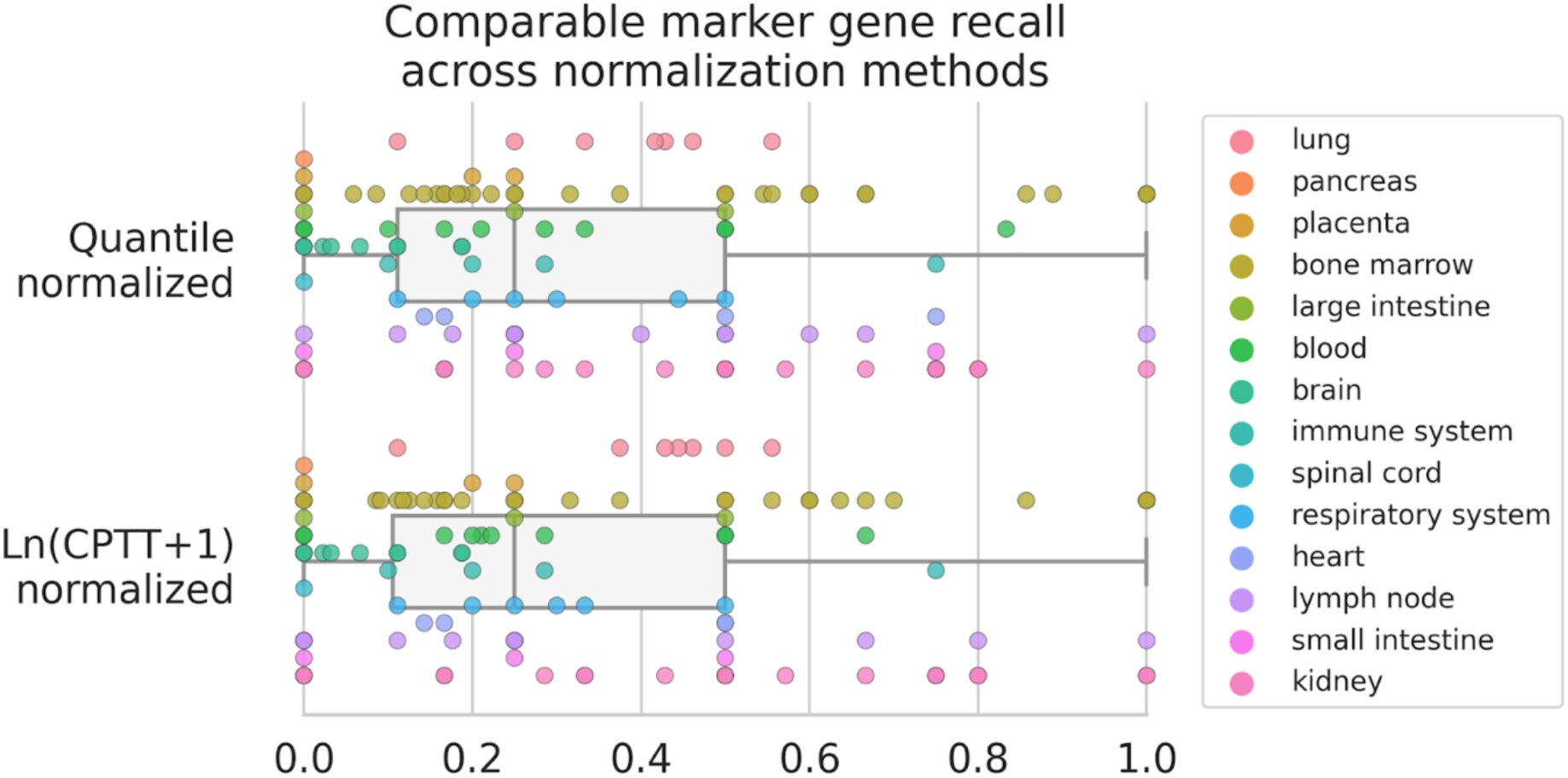
Recall of Marker Genes – Comparison between raw counts, quantile normalization and log transform normalization. Each point represents the sensitivity of marker gene recall for a specific cell type and tissue, as compared to canonical marker genes from HuBMAP.

**Figure 9.**
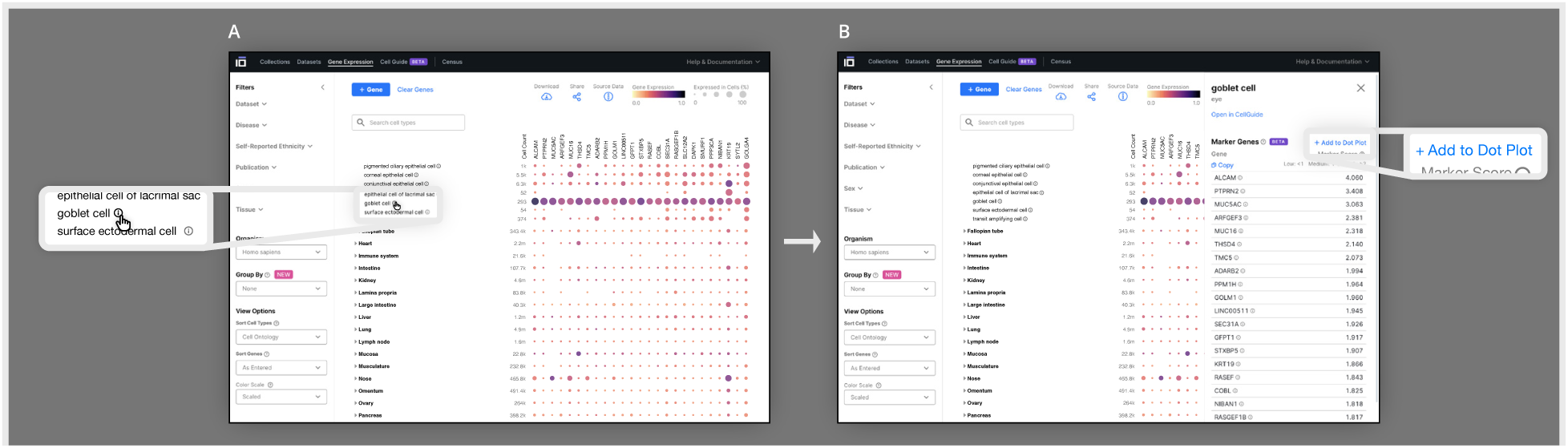
Visualizing marker genes across the data corpus. (A) From a Gene Expression heatmap generated from the log transform normalized data object, researchers can identify maker genes for any given cell type present in the normalized data object by clicking on the info button next to the cell type name on the heatmap. For each cell, its top 25 marker genes (marker score > 0.5) are calculated using Welch’s *t*-test to compare the gene expressions in the selected cell type to each other cell type in the tissue that is present in the data corpus. All marker genes for a given cell type can be quickly added to the heat map by selecting Add to Dot Plot.

**Figure 10.**
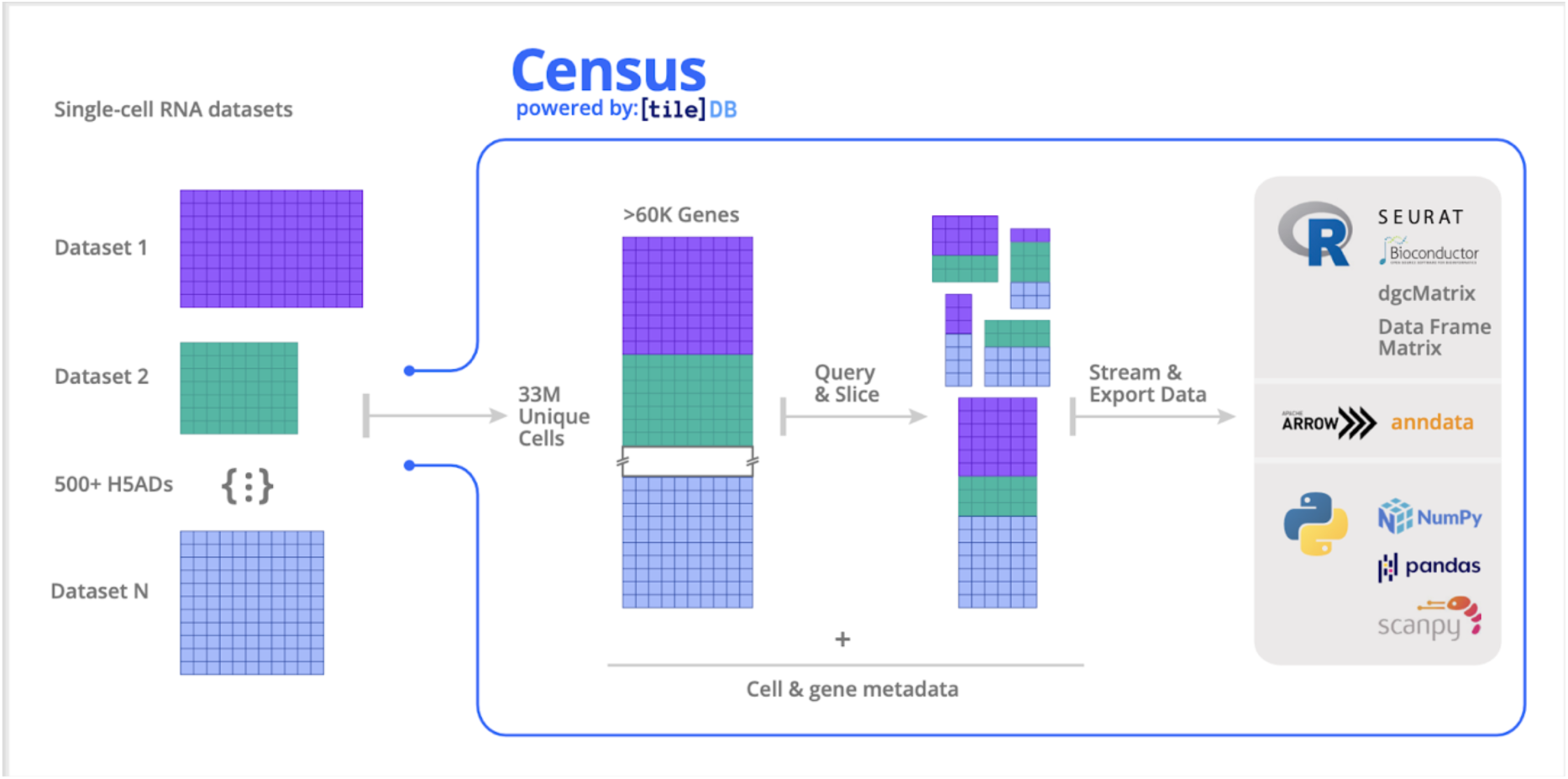
Overview of Census framework. Census is built upon the TileDB-SOMA framework to enable computational scientists to execute complex and specific queries across over 33 million cell measurements compiled from 500+ datasets spanning human and mouse organs available in CellxGene using Census. Leveraging out-of-core processing, SOMA provides the API and data model to facilitate the storage, retrieval, and analysis of over 500 datasets exceeding memory capacity. The standardized schema required by the data portal enables users to effortlessly query and export any segment of the extensive 33+ million cell dataset for in-depth analysis using Python and R.

### Census provides efficient programmatic access to CellxGene data for flexible exploration and reanalysis

A central goal of the CellxGene effort is to ensure robust programmatic access to the data for reuse and atlas-level analysis and modeling. To support this, we developed a global view of the data, termed Census, as a service composed of a cloud-hosted data object hosted at Amazon’s Open Data registry along with Python and R packages for efficient computing and access to non-spatial transcriptomic data available in CellxGene (see Methods).

As evidenced by the magnitude of the data in CellxGene, the single-cell community is now facing a challenge of data access and analysis at scale. Many tools that are widely used for analysis, or even meta-analysis, are not designed to operate on datasets of this scale.^40^ There are a myriad of existing computational analytical tools and methods available to the community, ^41,42,43,44^ many of which have made significant advancements towards increasing scalability around data access and analysis. For example, for R users, Seurat V5^45^ introduced a new paradigm to perform memory– and compute-costly processes on subsets of a dataset, while performing light-weight processes on full datasets. Similarly, for Python users AnnData^34^ introduced on-disk representations of data while incorporating state-of-the-art parallel computing via Zarr^46^ and Dask^47^ respectively. While these tools are increasing the scale to which computational biologists can access and analyze data, many of these solutions are still limited to only millions of cells, and more importantly, lack of interoperability between R and Python. To fill this gap, and in particular, to allow interoperable and scalable data access beyond 10s and 100s of millions of cells, we co-developed TileDB-SOMA with TileDB (**table S1**, see Methods). This API is intentionally designed for data access at scale while maintaining interoperability with the existing computational tools in the single-cell ecosystem. TileDB-SOMA was then used to build Census – a large data object hosted in the cloud with all RNA non-spatial transcriptomic data from CellxGene.

Through its cloud-based platform, Census offers efficient, low-latency access for larger-than-memory slices of CellxGene data, which users can access through Python and R APIs by performing queries based on cell or gene metadata. Census is interoperable across existing single-cell toolkits as query results can be exported to Seurat,^46^ AnnData,^34^ or SingleCellExperiment^44^ objects. Notably, Census offers interoperability with basic language structures: from Python, it can export data to PyArrow objects,^48^ SciPy sparse matrices,^49^ NumPy arrays,^50^ and Pandas data frames.^51^ From R, Census can export data to R Arrow objects, sparse matrices (via the Matrix package), and standard data frames and dense matrices.

Importantly, TileDB-SOMA allows Census to interact with CellxGene data in an out-of-core fashion via its iterable-based streaming capabilities (see Methods). Concretely, this means that data can be queried, processed, and analyzed in data chunks of fixed size. Algorithms can then be adapted to work on an incremental basis; particularly any computation that can operate independently on per-cell or per-gene data can be easily implemented in this manner.

Computations that require random access to the full query result or multiple passes through the data can be redesigned as an on-line algorithm.^52^ In Census, we have already deployed out-of-core implementations for some commonly used computations in single-cell, including incremental mean and variance calculation, and the ability to identify highly variable genes, all of which can now be executed in a regular 8GB memory laptop across 33 million cells and more than 60 thousand genes. These out-of-core functionalities are available in the Census Python package (**table S1**).

Modeling of single-cell data at scale is an important aspect of Census given its demonstrated uses for *in silico* experimentation,^53,54^ data integration and annotation,^55,56^ cell state prediction,^57,58^ and clinical applications.^59,60,61^ PyTorch^62^ is one of the most popular machine learning frameworks in single-cell, with notable models built using the PyTorch library including scvi-tools models, Geneformer,^63^ and scGPT.^64^ To allow existing and new machine learning models to be trained on Census-scale data, we implemented PyTorch iterable data loaders that work natively with Census via TileDB-SOMA. Leveraging the modular design of the PyTorch libraries, the Census data loaders can be easily utilized in new and existing model training pipelines, allowing models to be trained on Census-scale data using readily available computing resources.

To be included in Census, datasets must be generated from non-spatial RNA technologies, contain cells from human or mouse, provide raw counts, and utilize only standardized cell and gene metadata as described in the CellxGene dataset schema description above. More details on current inclusion criteria can be found within Census Documentation (**table S1**).

## Discussion

CellxGene has quickly demonstrated its potential for novel biological insights. Researchers using the platform recently uncovered the mechanism by which aging can drive B-cell lymphoma^65^ and the identified signaling gene sets involved in small cell lung cancer.^66^ The standardized data from CellxGene has enabled the development of new computational tools including UniCell: Deconvolve Base (UCDBase), a pre-trained, interpretable, deep learning model to deconvolve cell type fractions and predict cell identity across transcriptomic datasets;^67^ scTab, a cell type prediction model;^68^ and CSeQTL, a tool for mapping cell type-specific gene expression quantitative trait loci.^69^ Cross-consortia collaboration has been enabled by CellxGene through the development and refinement of the Human Reference Atlas, as well as other efforts to unify terminology and data infrastructure across consortia.^70^ Notably, 15 international consortia have published data to CellxGene, utilizing the minimal standard schema to drive interoperability and ensure reusability and accessibility of data generation efforts.

This compilation of standardized single-cell transcriptomic datasets lends itself to large-scale analysis, including the visualization of gene expression across various tissues in the human body. Through the Gene Expression interface, researchers can now quickly identify computationally computed marker genes for over 600 cell types and begin to understand in what cell types a specific gene of interest is expressed. We acknowledge that this method is imperfect as it does not fully correct for batch effects. However, it is well suited for our use cases: normalization with log transformed scaled pseudocounts meets the scalability requirements for processing large datasets efficiently, allowing for the iterative addition of new cells without necessitating a full re-run of normalization processes. It also successfully stabilizes variance for visualization purposes while providing a complete count matrix of expression values (rather than low-dimensional representations). These findings collectively support the suitability of log transform normalization for the Gene Expression tool so long as users are aware of its shortcomings.

The ability to access and download aggregate data from CellxGene through Census now provides researchers with an unprecedented scale and resolution to understand the molecular processes involved in health and disease. This increase in scale is coupled with an increase in obstacles related to data analysis, such as missing or noisy data, differentiating between technical and biological variance, overfitting, and data complexity, making it difficult to draw useful insights. Deep learning and generative modeling provide an exciting opportunity to address many of these challenges.^62,63^ An important bottleneck that CellxGene aims to alleviate is the lack of ready-to-train standardized data. A recent demonstration of the power of CellxGene data is on its application to train scGPT, a generative pre-trained model, on 33M cells,^64^ which the authors used for cell-type annotation, multi-batch integration, multi-omic integration, genetic perturbation prediction, and gene network inference. Other forms of global representations that utilize CellxGene are delivering value for scientists seeking to contextualize their data against large reference collections.^71^ Given the rapid and exciting advances in the field of Machine Learning and Artificial Intelligence, we anticipate that many additional applications of Large Language Models (LLMs) and modeling approaches will be powered by Census and the CellxGene suite of tools. Here we have presented the foundation of data aggregation, curation, and access that enables universal views of data but also lays a path for future modeling to capture the collective power presented by cell atlases.

As the single-cell biology field continues to evolve, it is critical that data portals and resources provide a consistent, reliable framework, while also adapting to the emerging technologies and the trajectory of the field. To this end, ongoing work includes updating all CellxGene features (Gene Expression, Marker Genes, Census) to present data that is derived from the same versioned snapshot of Portal data. Additionally, there is a need to further validate the normalization methods used across the corpus. Importantly, the field has acknowledged the importance of spatial information in the context of single-cell biology,^72,73^ and additional work is planned to support these efforts and technologies.

In summary, the CellxGene platform not only enables data sharing and data reuse through its standardized schema and curation pipeline but allows researchers across expertise to access and visualize the largest single-cell dataset to date, with over 50 million cells across 252 tissues. Critically, the release of the Census has unlocked a new potential for biological insights at scale and provides a necessary bridge between single-cell biology and the machine learning community, already signaling exciting biological and translational impact.

## Methods

### Performant visualization and analysis for up to 4M cells via Explorer

To achieve unparalleled efficiency, Explorer is engineered to incorporate numerous state-of-the-art tools and protocols. At the heart of its memory efficiency lies the use of TileDB Embedded (https://tiledb.com/products/tiledb-embedded/) as its data storage engine, a powerful, out-of-core solution for managing massive multi-dimensional array data. It excels at storing and accessing large and sparse datasets, making it an ideal choice for handling single-cell RNA sequencing data.

We utilized FlatBuffers to compress and quickly transfer large chunks of cell metadata and gene expressions to clients.

Categorical metadata is integer-encoded, and gene expressions are digitized to maximize compressibility.

To optimize the performance of differential expression, we engineered a custom implementation of Roaring Bitmaps, an efficient encoding method used for compressing postings lists, which contain cell indices in user requests. This algorithm and numerous other optimizations in the backend allow for the interactive comparison of populations of up to 1.5 million cells in under 60 seconds.

Finally, we built a custom renderer in WebGL, a JavaScript API for rendering high-performance graphics, to efficiently render millions of points in the cell embeddings and respond with minimal latency to user interaction.

### Data processing and Normalization for Gene Expression

#### Removal of Duplicate Cells

Some data on CellxGene is duplicated due to independent submissions, for example meta-analysis vs original data. All data submitted on Discover is curated to indicate whether any cell is the primary data. Only cells demarcated as primary data are included in the processing steps below.

#### Removal of Low Coverage Cells

Any cell that has less than 500 genes expressed is excluded, which filters out about 8% of all data and does not eliminate any cell type in its entirety. This filter enables more consistent quantile vectors used for the normalization step.

#### Removal of Cells Based on Sequencing Assay

Only cells from sequencing assays that measure gene expression and don’t require gene-length normalization are included (**Table 1**).

**Table 1.** Sequencing assays included in Gene Expression normalized data object.

| Assay | EFO ontology term ID |
| --- | --- |
| sci-RNA-seq | EFO:0010550 |
| 10x 3' v1 | EFO:0009901 |
| 10x 5' v1 | EFO:0011025 |
| 10x 3' v2 | EFO:0009899 |
| 10x 5' v2 | EFO:0009900 |
| 10x 3' v3 | EFO:0009922 |
| 10x 3' transcription profiling | EFO:0030003 |
| 10x 5' transcription profiling | EFO:0030004 |
| 10x technology | EFO:0008995 |
| Seq-Well | EFO:0008919 |
| Drop-seq | EFO:0008722 |
| CEL-seq2 | EFO:0010010 |

#### Data Normalization

Read counts are normalized using the ln(CPM+1) transformation of raw counts, where CPTT is Counts Per Ten Thousand.

Normalized matrices from multiple datasets of the same tissue are concatenated along the gene axis.

#### Removal of Noisy Ultra-low Expression Values

After applying normalization, any gene/cell combination counts less or equal than 2 are set to missing data. This allows for removal of noise due to ultra-lowly expressed genes and provides a cleaner visualization.

#### Summarization of Data in Heatmap

For each gene/cell type combination, the average ln(CPM+1)-normalized gene expression among genes that have a non-zero value are visualized by the dot color. The percentage of cells of any given cell type that express the gene of interest is visualized by the heatmap dot size, with the absolute number of cells expressing the gene found in parentheses. These numbers are calculated after the removal of low coverage cells.

#### Expression and cell count rollup across descendants in the cell ontology

The cell ontology is a hierarchical tree structure that represents the relationship between cell types. For example, the cell type “B cell” is a descendant of the cell type “lymphocyte”. For a particular cell type, the cell ontology is used to sum up the expression values and cell counts of cells labeled as that cell type as well as those labeled as its descendants. In the aforementioned example, the average expression of “lymphocyte” would include “B cells” and all its other descendants.

This rollup operation accounts for the fact that different datasets may have labeled their cells with different levels of granularity. It provides a more robust measure of the average expression of low-granularity cell type terms, such as “secretory cell” or “lymphocyte”.

### Data-generated marker genes

For each of the cell types available in the data corpus, we use a Welch’s *t*-test on the normalized values to compare the average expression of each gene in the cell type of interest against each other cell type in the same tissue. For each gene, we can take the 10th percentile of effect sizes across all these cell type comparisons as the reported effect size. However, for small numbers of comparisons, the 10th percentile can be a noisy metric. To improve its robustness, we bootstrap the distribution of effect sizes, taking the 10th percentile of each replicate, and averaging all replicates to get the final effect size. We return the 25 genes with the top effect sizes.

It is important to note that some methodological decisions were made to balance accuracy with efficiency and scalability. For example, we use a *t*-test to perform differential expression, which is a simple and fast test. However, it may not be as accurate as more sophisticated (and computationally intensive) statistical tests and differential expression values were calculated using normalized and log-transformed values instead of raw counts. Applying secondary filters to the data (like disease, ethnicity, etc.) does not affect the results. Enabling dynamic calculation of marker genes for arbitrary populations of cells in arbitrary subsets of the data may be a direction for future development.

### Biological validation of CellxGene normalization methods

#### 1. Batch Effects Analyses

*Note that the batch effect analyses used a scaling factor of 1,000,000 – rather than 10,000 – for the log transformation of scaled pseudocounts normalization. a*. Log fold changes of normalized expression values across covariates We normalized the data, stratified by covariate values, and then for each gene expressed in a particular cell, we computed the average expression value for each covariate value (e.g., each dataset, each assay, etc). In order to be consistent with the way averages are computed in the Gene Expression Application, we only considered non-zero gene expression values. We then calculated the log(fold change) between each pair of means per covariate. If a particular cell has a total of nk values for covariate *k*, we take the log fold changes between (nk choose 2) pairs. For example, if a type I pneumocyte in lung is present across 8 different datasets, we are are computing the pairwise log fold changes between a total of (8 choose 2) = 28 pairs (dataset1/dataset2, dataset1/dataset3, …, dataset7/dataset8, etc). We take the absolute value of the log(fold change). Each pair of covariate values is only considered once (e.g. if the pair dataset1/dataset2 is included, the pair dataset2/dataset1 is not). We assessed dataset_id and assay as potential covariates. The reported standard deviations were calculated using the formula for average standard deviation for k groups of unequal size by taking the square root of the sum of variances of each individual cell type under a different covariate, weighted by the sample size divided by the total number of observations:

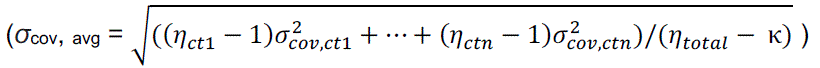

where n_ct1_, n_ct2_, …, n_ctn_ are the total number of samples for each cell type and σ*^2^_cov,ct_* are the variances of these cell types in a particular covariate

b. ANOVA on average normalized gene expression values across covariates We computed a one-way repeated measures ANOVA treating the average gene expression value of a particular gene as the dependent variable and the covariate as the independent variable. Average gene expression values were computed by excluding normalized gene expression values of 0. We looked at five different cell types and compared values across marker genes and housekeeping genes. Since there are genes that don’t appear across all covariate values, the system design was unbalanced. Our null hypothesis H0 was that there is no significant difference between the average normalized gene expression values across covariate values. For most cell types, the p_values obtained were greater than 0.05, which means that we did not have sufficient evidence to reject the null hypothesis. We used the pingouin package to compute the scores. We looked at Dataset ID and Sequencing Assay as potential covariates.

#### 2. Marker Gene Sensitivity

We selected 12 cell types across 6 tissues which had both canonical marker genes available in HuBMAP and raw count data available in CellxGene: Gene Expression data corpus. We then normalized the raw counts for each tissue using either ln(CPTT+1) or quantile normalization. Using these normalized values, we then computed marker genes for each of these cell types using the methods described above; however, we used a student’s t-test instead of Welch’s simply based on the availability of this method in scanpy rather than requiring the full production pipeline for the validation study.

### TileDB-SOMA development for Census

We worked in collaboration with TileDB to develop a technology for efficient and scalable single-cell data handling. Our efforts resulted in the abstract API specification, SOMA (“Stack of Matrices, Annotated”) (https://github.com/single-cell-data/SOMA) and its Python and R implementations via TileDB-SOMA (https://github.com/single-cell-data/TileDB-SOMA).

TileDB-SOMA was then used as the foundation to build Census for efficient programmatic access to CellxGene data. SOMA provides an API and data model for single-cell data to store and access larger-than-memory datasets by providing query-ready data management for reading and writing at low latency and cloud scale.

The data model behind the SOMA specification is flexible and extensible, and it is inspired by existing single-cell data formats, notably AnnData. It can accommodate multiple measurements from derivative views (e.g., spatial and non-spatial data) and embeddings of sparse and dense data, along with both observation (e.g., cells) and feature (e.g. genes) axis annotations.

Importantly the flexibility of the SOMA data model allows for representations beyond the single-dataset paradigm and enables managing single-cell data from multiple modalities (e.g., RNA, spatial, epigenomics) across joint or disjoint observations.

SOMA’s data model provides fundamental building blocks that can be composed into an arbitrary structure for a particular use case. These building blocks include: 1) Collections that act as containers for other SOMA data types, 2) Experiments that are specialized collections to meaningfully group single-cell data measurements and allow for observation– and feature-based queries, 3) Measurements that are specialized collections of single-cell data from a shared molecular measurement (e.g. RNA or protein), 4) Data Frames that provide multi-column tables, and 5) Multi-dimensional Arrays (Sparse and Dense) to represent the multi-dimensional numeric data.

Leveraging this model, Census data is packaged into multiple Experiments, one per organism (*homo sapiens* and *mus musculus*, currently). Each experiment contains a Measurement for RNA data and an “obs” Data Frame for cell metadata and annotations. The Measurement contains a “var” Data Frame for gene metadata and a Collection of “X” (expression data) layers. Currently, two layers are provided: One for RNA transcript raw counts and one for library-size normalized counts (Figure 11). Census data is packaged into multiple Experiments, one per organism (homo sapiens and mus musculus, currently). Each experiment contains a Measurement for RNA data and an “obs” Data Frame for cell metadata and annotations. The Measurement contains a “var” Data Frame for gene metadata and a Collection of “X” (expression data) layers. Currently, two layers are provided: One for RNA transcript raw counts and one for library-size normalized counts. Finally, an additional top-level Collection of summary information Data Frames provide a profile of the contents of the Census data.

Fundamentally, the SOMA API’s design differentiates from existing technologies by allowing for and assuming out-of-core processing as the primary default use case. All query results are returned to the client via an iteration pattern that streams the data to the client incrementally, limiting the size of the data that must be handled at any given time to a fixed size “chunk”.

import CellxGene_census as cc

import tiledbsoma as soma

census = cc.open_soma()

experiment = census[“census_data”][“homo_sapiens”]

axis_query = soma.AxisQuery(value_filter=“tissue_general == ‘lung’”)

with experiment.axis_query(measurement_name=“RNA”,

obs_query=axis_query) as query:

\# Iterate over X data, returning “chunks” of PyArrow tables

for table in query.X(“raw”).tables():

\# PyArrow table can be converted to a Pandas DataFrame, e.g.

df = table.to_pandas()

\# Do something with df

print(df)

\# Iterate over X data, returning “chunks” of COO Matrices

for coo_matrix in query.X(“raw”).coos():

\# Do something with coo matrix

print(coo_matrix)

In many cases, algorithms can be easily adapted to work with this incremental data access pattern. In particular, any computation that can operate independently on per-cell data should easily accommodate the SOMA API. However, computations that require random-access to the full query result or multiple passes through the data may need to be redesigned as an on-line algorithm. For example, computation of variance can be performed using Welford’s online algorithm.^74^

The SOMA specification is currently implemented by TileDB-SOMA, a Python and R client library that leverages TileDB (https://tiledb.com/) as its underlying database technology. By providing SOMA as a *specification* and TileDB-SOMA as one possible *implementation*, we aim to encourage the single-cell community to adopt the specification more broadly.

TileDB is an embedded database that enables a server-less architecture for accessing data stored in the cloud. TileDB’s first-class support for cloud data storage means that it is highly optimized to minimize network latency and accomplishes this via use of indexing and compression techniques. As the Census is currently hosted on AWS S3, access to its data is effectively limited only by a user’s network bandwidth and compute resources. The TileDB querying engine supports efficient multi-dimensional slicing (i.e., fast reads), which naturally supports querying for Census data over the obs and var axes. CZI’s single-cell team had previously employed TileDB technology for other CellxGene applications, including Explorer and Gene Expression.

## Availability and contact

All source code for CZ CellxGene tools can be found at https://github.com/orgs/chanzuckerberg/repositories?&q=cellxgene. CellxGene data can be accessed at https://CellxGene.cziscience.com/collections. The data and code used to assess the normalization methods behind CellxGene: Gene Expression can be found at https://github.com/chanzuckerberg/cellxgene-manuscript-2023. For additional questions, please contact.

## Acknowledgements

CellxGene has been a community effort. We would like to thank all the data curators, engineers, and subject matter experts who have consulted on or contributed to the CellxGene resource, including: Madison Dunitz, Ronen Kalo, Marcus Kinsella, Vlad Kislev, Ed Lein, Malte Lucken, David Osumi-Sutherland, Angela Pisco, Sara Simmonds, Kyle Travaglini, Galabina Yordanova. Additionally, we would like to thank Jauslyn Robbins for team administrative support, Giulia Notari for figure design, Alex Lokshin, Jake Heath, and the CZI Central Infra team for their technical assistance and support, the CZI Human Resources team for staffing support, Jae Lee and the entire CZI Central Trust Team for their support and guidance with data privacy, and Nicki Ghafari Saravi and the CZI Science Communications team. We would also like to acknowledge and thank all researchers, labs, and consortium members who have contributed data to CellxGene.

