## Supplemental Materials for "CZ CELL×GENE Discover: A single-cell data platform for scalable exploration, analysis and modeling of aggregated data"

### Supplemental Figures & Methods

**Supplemental Table 1**. CellxGene documentation and resource links

| Description | Link |
|---|---|
| CellxGene Schema | https://github.com/chanzuckerberg/single-cell-curation/tree/main/schema |
| CellxGene schema changelog | https://github.com/chanzuckerberg/single-cell-curation/blob/main/schema/3.1.0/schema.md#appendix-a-changelog |
| CellxGene data eligibility criteria and submission process | https://cellxgene.cziscience.com/docs/032__Contribute%20and%20Publish%20Data |
| Gene Expression data processing & normalization documentation | https://cellxgene.cziscience.com/docs/04__Analyze%20Public%20Data/4_2__Gene%20Expression%20Documentation/4_2_3__Gene%20Expression%20Data%20Processing#data-normalization |
| Gene Expression source data | https://cellxgene.cziscience.com/docs/04__Analyze%20Public%20Data/4_2__Gene%20Expression%20Documentation/4_2_6__Gene%20Expression%20Source%20Data |
| TileDB-SOMA | https://github.com/single-cell-data/TileDB-SOMA |
| Census Python API | https://chanzuckerberg.github.io/cellxgene-census/python-api.html |
| Census data and schema | https://chanzuckerberg.github.io/cellxgene-census/cellxgene_census_docsite_schema.html#data-included-in-the-census |

**Supplemental Table 2.** Metadata fields and corresponding ontologies

| Metadata field | Ontology/Standards | Ontology link |
|---|---|---|
| organism (species) | NCBITaxon | http://obofoundry.org/ontology/ncbitaxon.html |
| development_stage (age) | HsapDv/MmusDv | https://obofoundry.org/ontology/hsapdv.html<br>https://obofoundry.org/ontology/mmusdv.html |
| is_primary_data | Boolean (true/false) | N/A |
| gene IDs | Ensembl | https://useast.ensembl.org/Help/View?id=285 |
| sex | PATO | https://obofoundry.org/ontology/pato.html |
| self_reported_ethnicity | HANCESTRO | https://obofoundry.org/ontology/hancestro.html |
| disease | MONDO/PATO | https://obofoundry.org/ontology/mondo.html<br>https://obofoundry.org/ontology/pato.html |
| tissue | UBERON | https://obofoundry.org/ontology/uberon.html |
| suspension_type | [cell,nucleus] | N/A |
| assay | EFO | https://www.ebi.ac.uk/efo/ |
| cell_type | CL | https://obofoundry.org/ontology/cl.html |

**Supplemental Table 3.** List of Markers and Housekeeping Genes Used for Normalization Analyses

| cell_type | tissue | marker genes | housekeeping genes |
|---|---|---|---|
| alveolar type 2 fibroblast cell | lung | LUM, SCARA5, FBLN1, DCN, COL1A2 | MALAT1, ACTB, GAPDH, UBC, SDHA |
| syncytiotrophoblast cell | placenta | KRT7, EGFR, PSG4, CSH1 | MALAT1, ACTB, GAPDH, UBC |
| T cell | kidney | CD3G, CD40LG, CD3D, CD247, CD96 | MALAT1, ACTB, GAPDH, UBC, SDHA |
| plasma cell | lymph node | IRF4, SDC1, PRDM1, CD38 | MALAT1, ACTB, GAPDH, UBC |
| plasma cell | bone marrow | IGHG1, IGHA1, SEC11C, SDC1, DERL3 | MALAT1, ACTB, GAPDH, UBC, SDHA |

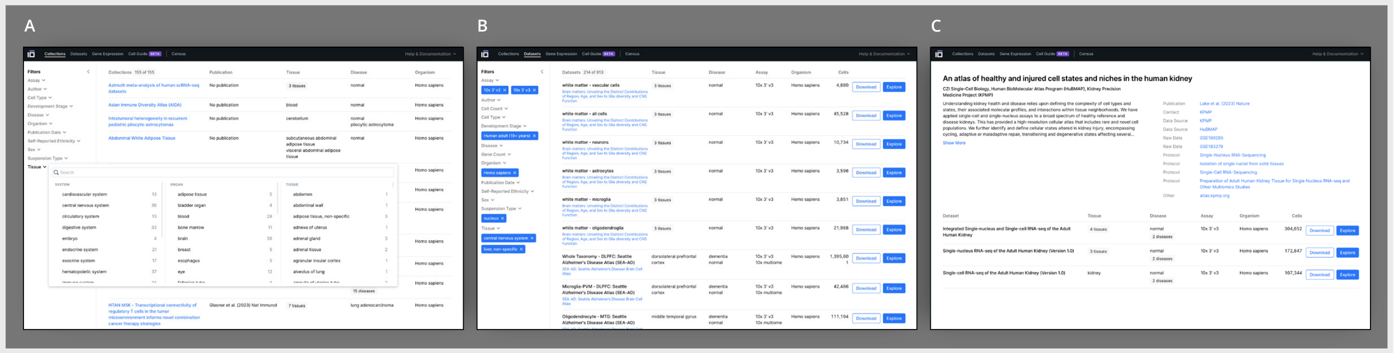


**Supplemental Figure 1**. (A,B) CellxGene User Interface (UI) allows for researchers to quickly filter and sort through datasets and collections based on collection metadata. (C) Each collection has an associated collections page with details about the author, study, datasets, and other collection metadata. From the collections page, researchers can easily explore any dataset that is part of the collection.

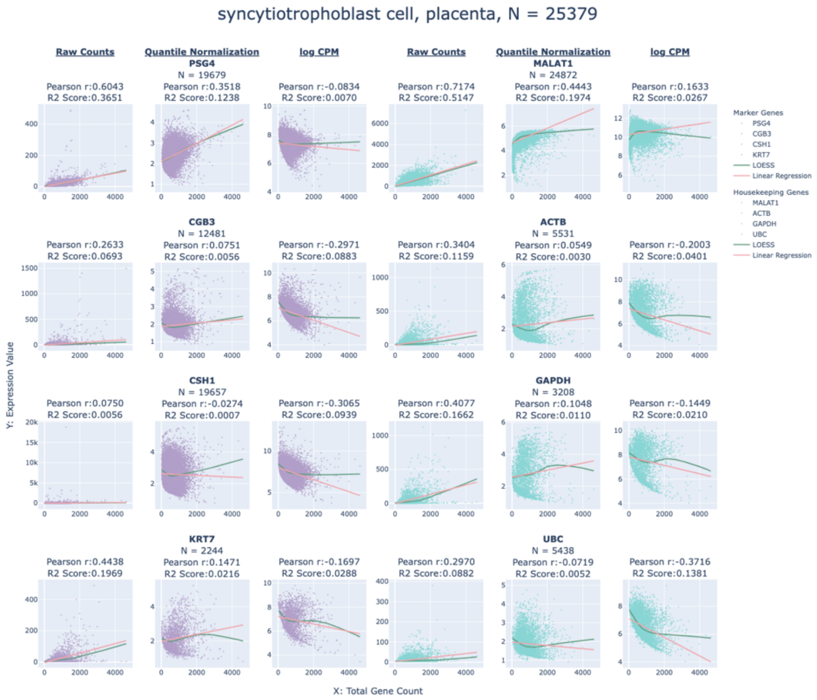


**Supplemental Figure 2**. **Gene Expression Value as a function of Total Gene Count for raw counts, quantile and log CPM normalizations** for *syncytiotrophoblast cells in placenta* (N = 25379 observations). The X axis represents the total number of genes expressed in a cell, and the Y axis represents the expression values (raw or normalized). N below each gene represents the number of non-zero gene expressions observed for that gene in the cell type. Markers were retrieved from HuBMAP.

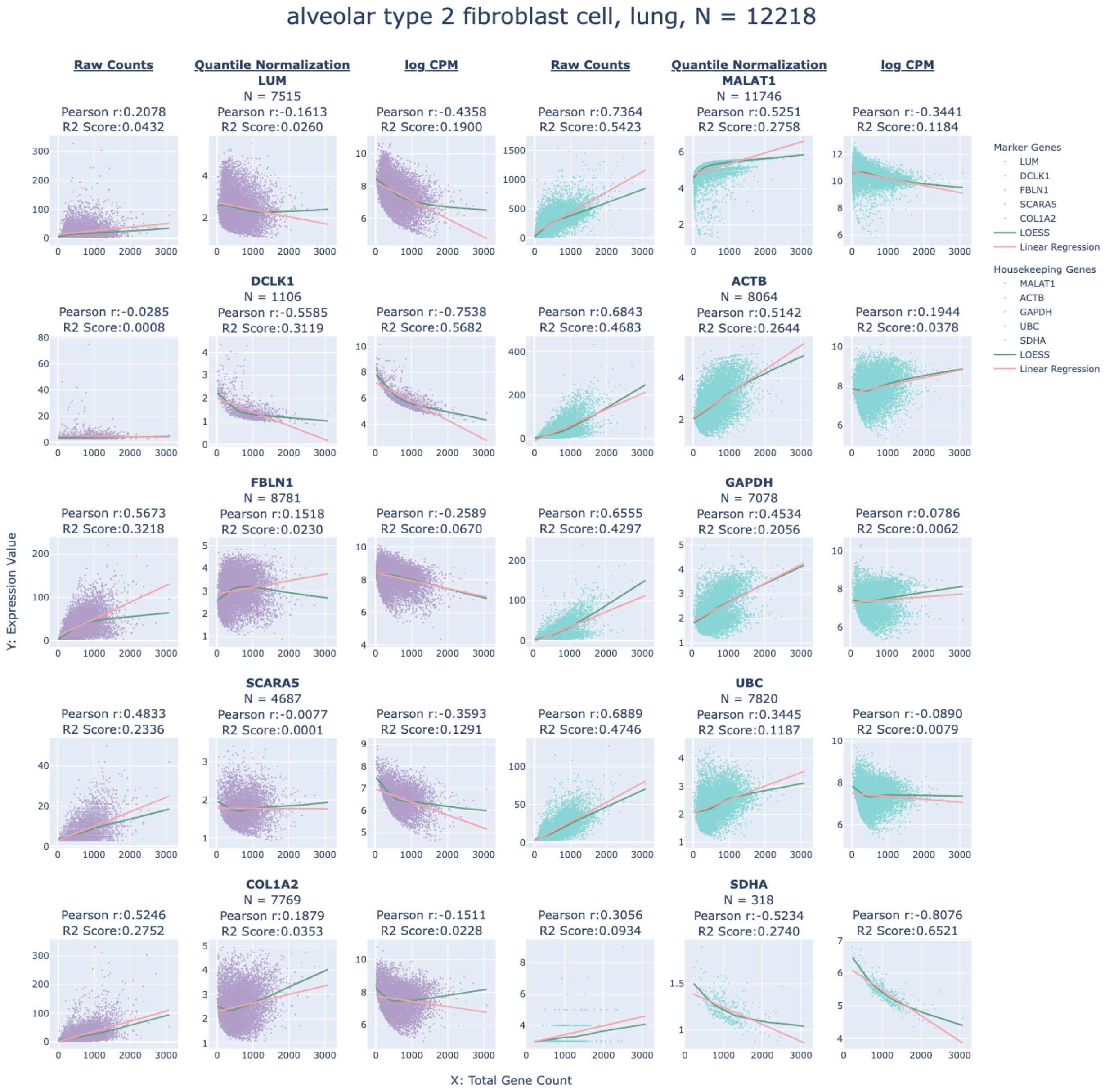


**Supplemental Figure 3**. **Gene Expression Value as a function of Total Gene Count for raw counts, quantile and log transform normalizations** for *alveolar type 2 fibroblast cells in lung* (N = 12218 observations). The X axis represents the total number of genes expressed in a cell, and the Y axis represents the expression values (raw or normalized). N below each gene represents the number of non-zero gene expressions observed for that gene in the cell type. Markers were retrieved from HuBMAP. The scaling factor for the log transform normalization used in this analysis was 1,000,000.

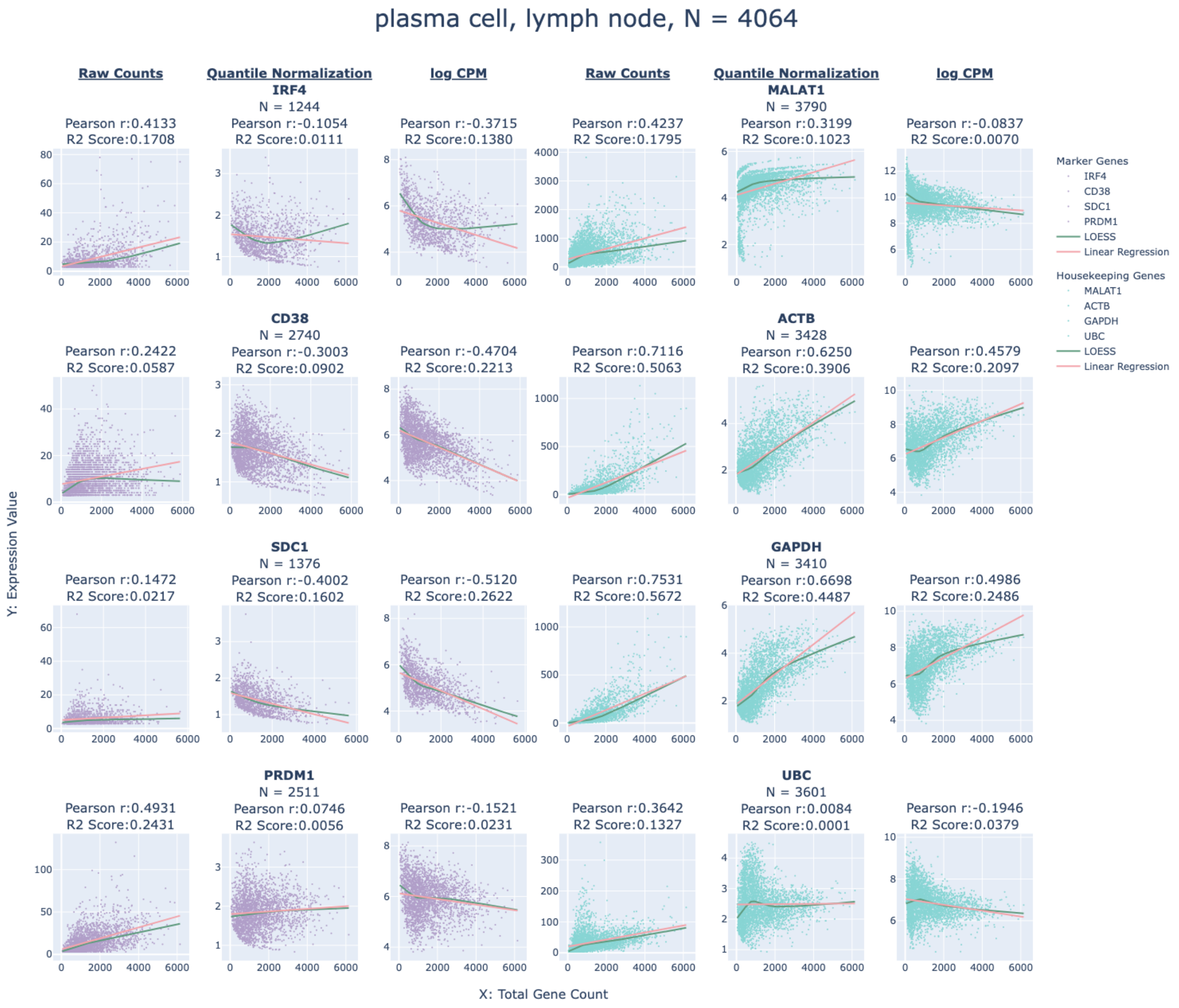


**Supplemental Figure 4. Gene Expression Value as a function of Total Gene Count for raw counts, quantile and log CPM normalizations** for *plasma cell in lymph node* (N = 4064 observations). The X axis represents the total number of genes expressed in a cell, and the Y axis represents the expression values (raw or normalized). N below each gene represents the number of non-zero gene expressions observed for that gene in the cell type. Markers were retrieved from HuBMAP. The scaling factor for the log transform normalization used in this analysis was 1,000,000.

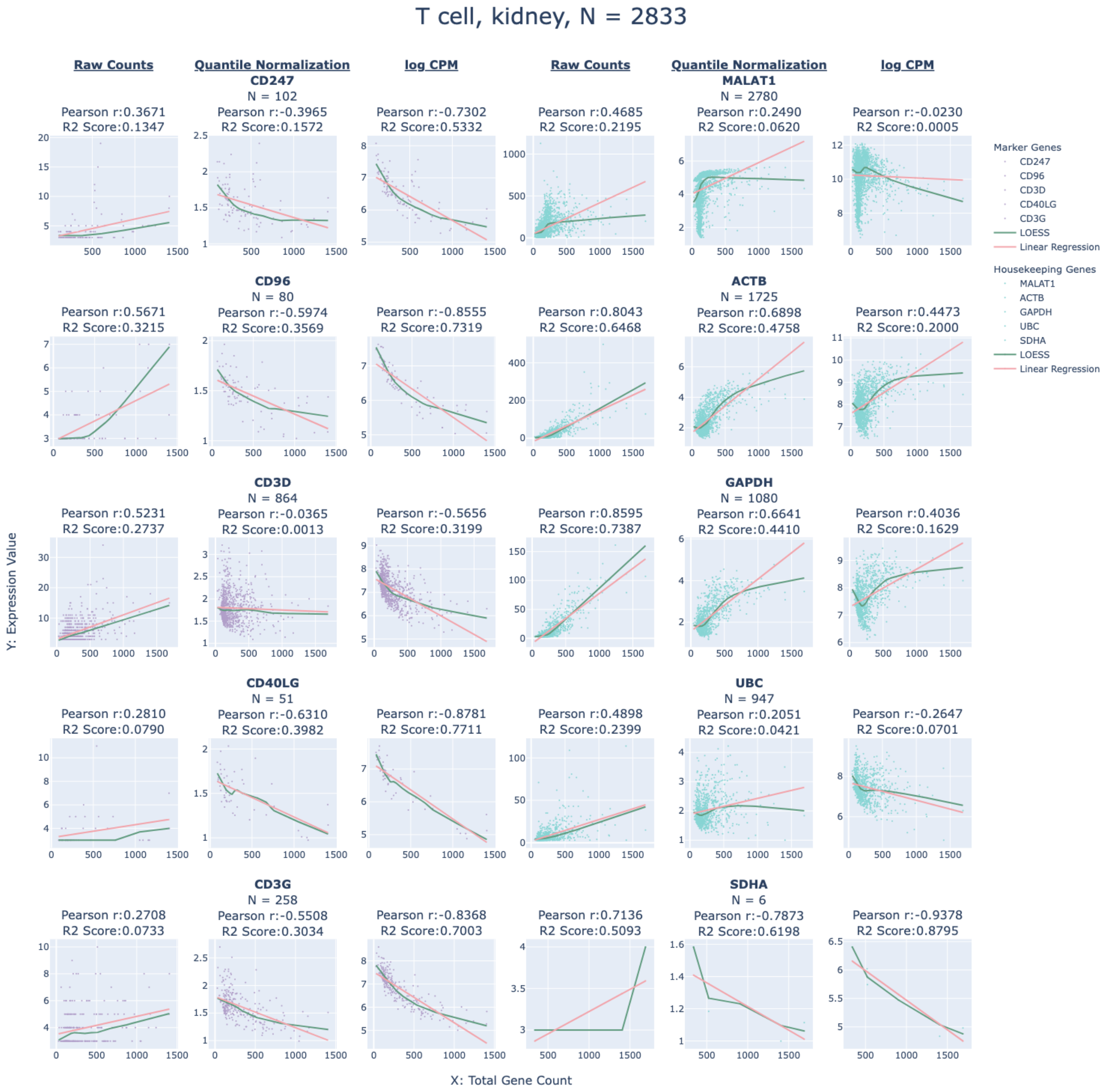


**Supplemental Figure 5**. **Gene Expression Value as a function of Total Gene Count for raw counts, quantile and log transform normalizations** for *T cells in the kidney* (N = 2833 observations). The X axis represents the total number of genes expressed in a cell, and the Y axis represents the expression values (raw or normalized). N below each gene represents the number of non-zero gene expressions observed for that gene in the cell type. Markers were retrieved from HuBMAP. The scaling factor for the log transform normalization used in this analysis was 1,000,000.

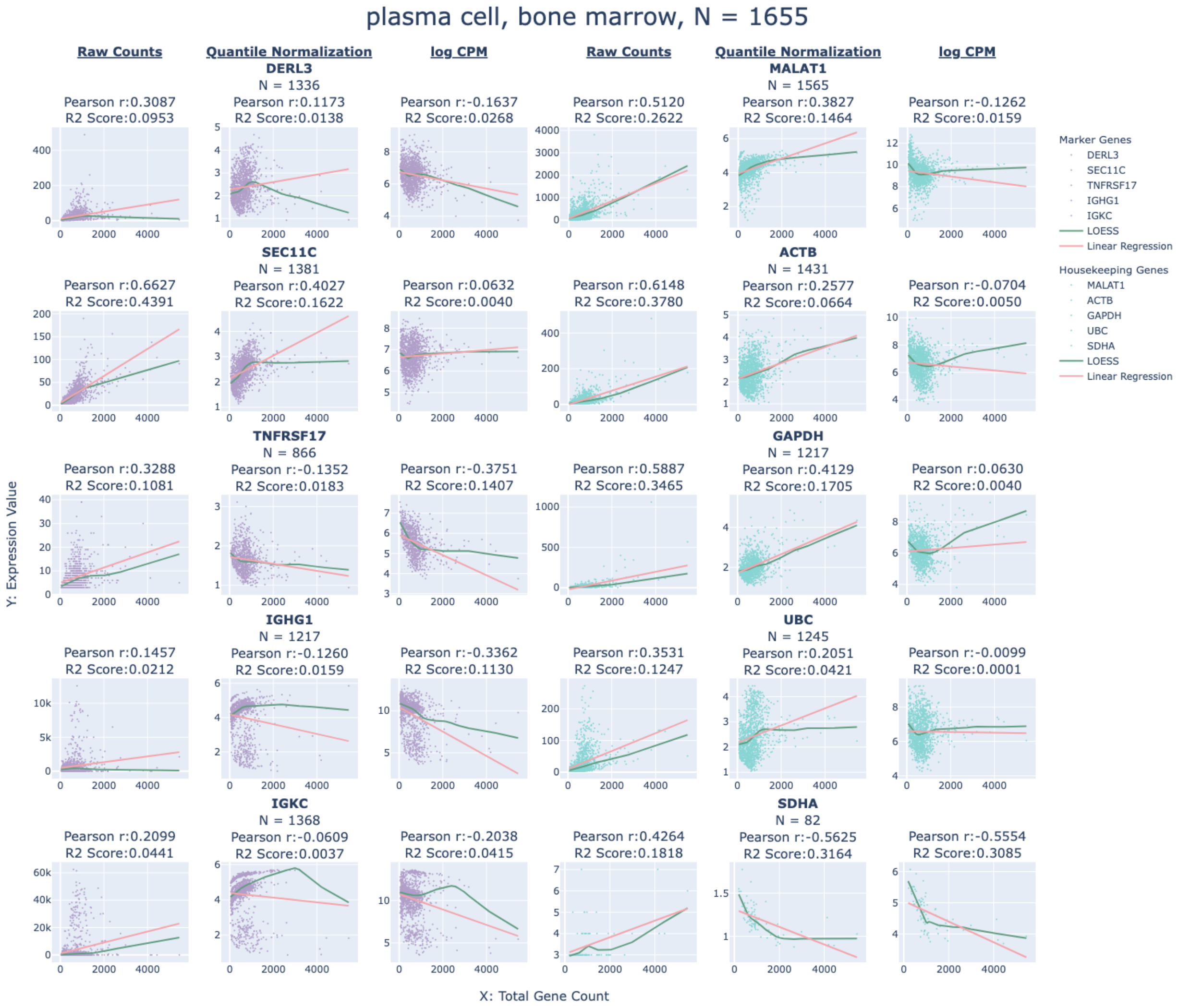


**Supplemental Figure 6**. **Gene Expression Value as a function of Total Gene Count for raw counts, quantile and log transform normalizations** for *plasma cell in bone marrow* (N = 1655 observations). The X axis represents the total number of genes expressed in a cell, and the Y axis represents the expression values (raw or normalized). N below each gene represents the number of non-zero gene expressions observed for that gene in the cell type. Markers were retrieved from HuBMAP. The scaling factor for the log transform normalization used in this analysis was 1,000,000.

### Supplemental methods

##### Influence of raw counts distribution on normalized values

We plotted the number of genes expressed per cell against the normalized gene expression values for a number of markers and housekeeping genes for four different cell types in five different tissues: alveolar type 2 fibroblast in lung, syncytiotrophoblast cell in placenta, T cell in kidney, and plasma cell in lymph node and bone marrow. We chose cells for which we could retrieve marker genes from HuBMAP. We used Pearson correlation (from pandas package) to assess the strength of the linear relationship between normalized gene expression and the number of genes expressed in a cell type. We used the $R^2$ score from a fitted Linear Regression model (from scikit-learn) to assess the contribution of variance explained by the total number of genes expressed in a cell in determining normalized gene expression values. We did this analysis for three different methods: raw counts, quantile normalization and log CPM normalization. The analyses excluded observations where a gene expression value was 0.

For each cell type, we plotted the distributions for a list of markers and housekeeping genes. The full list of markers, obtained from HuBMAP, and housekeeping genes are available in (table S3). We averaged the $R^2$ scores over marker genes and housekeeping genes across all cell types and reported the aggregated values **(figs. S2-6)**.
